# Hindlimb Muscle Representations in Mouse Motor Cortex Defined by Viral Tracing

**DOI:** 10.1101/2022.05.19.492320

**Authors:** Lauren Maurer, Maia Brown, Tamandeep Saggi, Christi L. Kolarcik

**Author notes:** Corresponding Author Name and Email Address: Christi L. Kolarcik.

## Abstract

Descending pathways from the cortex to the spinal cord are involved in the control of natural movement. Although mice are widely used in the study of the neurobiology of movement and as models of neurodegenerative disease, an understanding of motor cortical organization is lacking, particularly for hindlimb muscles. In this study, we used the retrograde transneuronal transport of rabies virus to compare the organization of descending cortical projections to fast- and slow-twitch hindlimb muscles surrounding the ankle joint in mice. Although the initial stage of virus transport from the soleus muscle (predominantly slow-twitch) appeared to be more rapid than that associated with the tibialis anterior muscle (predominantly fast-twitch), the rate of further transport of the virus to cortical projection neurons in layer V was equivalent for the two injected muscles. After appropriate survival times, dense concentrations of layer V projection neurons were identified in three cortical areas: the primary motor cortex (M1), secondary motor cortex (M2), and primary somatosensory cortex (S1). The origin of the cortical projections to each of the two injected muscles overlapped almost entirely within these cortical areas. This organization suggests that cortical projection neurons maintain a high degree of specificity, even when they are involved in controlling muscles with different functional roles (fast-versus slow-twitch, extensor versus flexor). Our results represent an important addition to the understanding of the mouse motor system and lay the foundation for future studies investigating the mechanisms underlying motor system dysfunction and degeneration in diseases such as amyotrophic lateral sclerosis and spinal muscular atrophy.

## 1. Introduction

Descending motor pathways from the cerebral cortex to the spinal cord mediate the movement of muscles throughout the body for behaviors ranging from basic locomotion to dexterous object manipulation. In rodents, these cortical projections originate in layer V and connect to brainstem nuclei and/or spinal cord interneurons. In humans and some non-human primates, cortical projections originating in layer V can directly impact the activity of motor neurons, and prior work indicates that these direct connections mediate highly-skilled motor behaviors in these species (1-3). Together, these direct and indirect pathways from the cerebral cortex determine how and when voluntary movement occurs. An understanding of these pathways is critical to identifying and targeting treatments for dysfunction associated with pathological conditions that affect the motor system.

Although the mouse is widely used to study the neurobiology of disease, the organization of the mouse motor system has not yet been fully delineated. Previous work by others has investigated the neural circuitry underlying forelimb muscles in mice, revealing a selective synaptic connectivity matrix between the cortex, brainstem, and spinal motor neurons that regulates forelimb movement (4, 5). However, the areas of the cortex and brainstem that are directly responsible for controlling the spinal cord motor neurons that innervate hindlimb muscles have not been fully characterized.

Another important consideration in understanding descending control over movement is the presence of different motor unit types. Muscles are composed of varying types of muscle fibers including fast fatigue-resistant (FR), fast fatiguable (FF), and slow-twitch (ST), and the motor neurons that innervate each muscle fiber type have different anatomical, physiological, and functional properties. Henneman’s size principle has established that small motor units and small motor neurons are recruited before large motor units and large motor neurons (6). These small motor units tend to be slow-twitch units that are recruited prior to fast fatigue-resistant and fast fatigable units. Differential loss of muscle fibers based on type has been observed in conditions ranging from muscular dystrophies to cancer cachexia [reviewed in (7)]. A preferential loss of fast-twitch muscle fibers has also been reported in rodent models of amyotrophic lateral sclerosis (ALS) (8-10) despite it being regarded as a disease of the motor neuron. A better understanding of differences in the cortical innervation of specific motor unit types is essential for addressing the neural mechanisms underlying these complex conditions.

In this study, we used trans-synaptic virus-based tracing to identify the origin of cortical control of motor neurons that innervate distinct hindlimb muscles in the mouse. We chose rabies virus (RV) to interrogate the motor circuitry because it is transported exclusively in the retrograde direction and trans-synaptically in a time-dependent manner (11); through the adjustment of survival time, retrograde transneuronal transport of RV enables identification of multiple nodes in a chain of synaptically-linked neurons (12). We investigated the innervation of two muscles with either predominantly slow-twitch muscle fibers (soleus, 50% type I) or predominantly fast-twitch muscle fibers (tibialis anterior, 90% type II) (8). We show that the motor neurons of these two hindlimb muscles were infected by retrograde transport of RV at different rates. Labeling of the soleus motor neuron pool occurred before that of the TA motor neuron pool, indicating differential uptake of RV at the level of muscle. However, following uptake of RV into the nervous system at the neuromuscular junction, the rate of retrograde transneuronal transport was not significantly different for soleus versus TA, and similar anatomical nodes in the spinal cord and brainstem were involved in mediating virus transport from both TA and soleus to layer V neurons in the cerebral cortex. Further, density-based analyses revealed that the cortical regions associated with the control of each hindlimb muscle exhibited considerable overlap. These results suggest that cortical neurons retain some differential specificity related to the muscle fiber types that innervate their target motor neurons.

## 2. Materials and Methods

### 2.1 Surgical Procedures

All surgical procedures were performed in accordance with those outlined by the Association for Assessment and Accreditation of Laboratory Animal Care (AAALAC) and the National Institutes of Health Guide for the Care and Use of Laboratory Animals. The experimental protocol was approved by the Institutional Animal Care and Use Committee as well as the Biosafety Committee of the University of Pittsburgh. Biosafety practices followed those outlined in Biosafety in Microbiological and Biomedical Laboratories (Department of Health and Human Services Publication No. 93-8395). Procedural details regarding the handling of virus and virus-infected animals have been outlined previously (13). Animals were housed in the facilities of the University of Pittsburgh, Division of Laboratory Animal Resources, and all efforts were made to minimize the number of animals used as well as their suffering.

All procedures were performed using aseptic techniques and under anesthesia. Adult male mice on the C57Bl/6 background (The Jackson Laboratory, Farmington, CT) weighing approximately 30 g were given buprenorphine (0.1 mg/kg) prior to and following surgery. Animals were anesthetized with 2.5% isoflurane in oxygen at approximately 150 mL/min and monitored closely throughout the procedure by observing changes in respiratory and heart rates. Animal temperature was maintained throughout the procedure using a heating pad, and ophthalmic ointment was applied to the eyes while under anesthesia.

Once anesthetized, animals were placed onto their left side and the right hindlimb shaved and then cleaned with betadine. The right hindlimb was draped with a sterile towel, and a longitudinal incision from approximately the knee to the ankle was made to expose either the tibialis anterior (TA) or the soleus muscle. Rabies virus (CVS-N2c; 1 × 10^9^ pfu/mL; supplied by M. Schnell, Thomas Jefferson University, Philadelphia, PA) was injected using a 30-gauge Hamilton syringe along the length of the muscle at a volume proportional to the weight of each muscle [approximately 3 µL for soleus and approximately 19 µL for TA based on muscle weights of 6.58 ± 2.09 mg and 42.17 ± 6.33 mg, respectively (14)]. Once injections were completed, sterile saline was used to flush the area surrounding the injection site. The skin was sutured, and the animals were returned to cages designed for the housing of virus-infected animals. Animals were singly-housed and given free access to food and water following surgery with environmental enrichment provided in the isolation suite throughout the survival period. Animals displayed no symptoms of rabies virus infection over the survival times utilized in this study. Current evidence suggests that rabies virus is transported transneuronally in all types of systems and across all types of synapses (11-13, 15).

### 2.2 Survival Period

The survival times varied from approximately 24 hours to approximately 93 hours. After the designated survival period, animals were deeply anesthetized with sodium pentobarbital (100 mg/kg) before being transcardially perfused first with 0.1 M phosphate buffer followed by either 10% phosphate buffered formalin or 4% paraformaldehyde. Following perfusion, the brain and spinal cord were removed and placed in fixative at 4°C for up to three days and then transferred to phospho-tris-azide. Brains were embedded in gelatin and serial coronal sections (50 µm) of the cerebral cortex and cerebellum were collected. Lumbar spinal cord blocks (from approximately T13 to S2) were embedded in gelatin and cross-sections (50 µm) were collected.

### 2.3 Histological Procedures

Every 10^th^ section of the brain and spinal cord was stained with cresyl violet to enable analysis of cytoarchitecture. Even-numbered sections of the brain and spinal cord were used to evaluate the presence of rabies-positive neurons. Immunohistochemical reactions on free-floating brain and spinal cord sections using the avidin-biotin peroxidase method (Vectastain, Vector Laboratories, Burlingame, CA) were used to identify rabies-infected neurons. These reactions utilized a mouse monoclonal antibody (31G10, used at a concentration of 0.1 µg/mL, supplied by M. Schnell, Thomas Jefferson University, Philadelphia, PA) incubated overnight at 4°C. After washing, biotinylated secondary antibody was added at a dilution of 1:500. The signal was then amplified using ABC Reagent with 3,3’-diaminobenzidine (DAB) as the chromagen. Reacted sections were mounted on gelatin-covered glass slides, air-dried, washed in distilled water, and coverslipped with Cytoseal.

For immunofluorescence, free-floating tissue sections were blocked in normal donkey serum in phosphate buffered saline (PBS). Primary antibodies were added for 48 hours at 4°C and included the mouse monoclonal antibody described above (31G10) (5) as well as commercially available cell type-specific markers including: choline acetyltransferase (ChAT; AB143; rabbit polyclonal; Millipore, Burlington, MA); COUP TF1-interacting protein 2 (CTIP2; rabbit polyclonal; Synaptic Systems, Goettingen, Germany); and microtubule associated protein 2 (MAP2; guinea pig polyclonal; Synaptic Systems). The appropriate fluorescence-conjugated secondary antibodies were added for 1-2 hours at room temperature at a dilution of 1:500. Hoechst 33342 (Invitrogen, Waltham, MA) was used as a nuclear counterstain according to the manufacturer’s instructions. Sections were mounted with SlowFade Gold (Invitrogen). Digital images were acquired with a fluorescence microscope (FluoView 1000, Olympus, Inc., Tokyo, Japan). Detection parameters were optimized for each fluorophore and consistent settings used for all images with image contrast and brightness adjusted in Adobe Photoshop (Adobe Systems Inc., San Jose, CA).

### 2.4 Neuronal Quantification

Spinal motor neurons were positively identified by labeling with RV (31G10 antibody) in every other section of the lumbar spinal cord. Counts of spinal motor neurons were performed in animals in which no spinal interneurons were labeled with RV; these counts were doubled to account for the sections that were not evaluated. In these animals, the virus-labeled motor neurons are reported as a percentage of the motor neuron pool (16, 17) (**Table 1**). Cortical neurons in layer V positive for RV were quantified from every other coronal section of the brain. Counts of neurons labeled in cortical layer V were performed in animals in which cortical labeling was confined to layer V. The same assessment was done for each hindlimb muscle at several survival times (**Table 2**).

**Table 1.** Quantification of spinal motor neuron labeling in animals with labeling restricted to first-order motor neurons following rabies virus (1 × 10^9^ pfu) injection. *Average number of total motor neurons based on spinal motor neuron pool of 142 for tibialis anterior and 41 for soleus (16, 17).

| | Survival time ($n$ = number of animals) | Percentage of the spinal motor neuron pool* labeled (average ± SD) |
| --- | --- | --- |
| **Tibialis anterior** |  |  |
| | 29 hours ($n$ = 3) | 3.3% ± 1.6% |
| | 36 hours ($n$ = 3) | 9.8% ± 3.7% |
| | 40 hours ($n$ = 4) | 12.7% ± 8.2% |
| | 44 hours ($n$ = 3) | 7.0% ± 5.1% |
| **Soleus** |  |  |
| | 24 hours ($n$ = 3) | 39.0% ± 25.4% |
| | 30 hours ($n$ = 4) | 56.0% ± 11.6% |
| | 36 hours ($n$ = 4) | 54.8% ± 14.0% |

**Table 2.** Quantification of layer V cortical neuron labeling in the hemisphere contralateral to the injection site in animals with labeling restricted to third-order layer V neurons following rabies virus (1 × 10^9^ pfu) injection.

| | Survival time ($n$ = number of animals) | Layer V neurons (average ± SD) |
| --- | --- | --- |
| **Tibialis anterior** |  |  |
| | 60 hours ($n$ = 2) | 1.5 ± 0.71 (*not included in density analyses*) |
| | 63 hours ($n$ = 1) | 6.0 ± 0.0 |
| | 69 hours ($n$ = 4) | 9.0 ± 8.5 |
| | 75 hours ($n$ = 2) | 86.5 ± 16.26 |
| | 84 hours ($n$ = 4) | 309.5 ± 251.84 |
| **Soleus** |  |  |
| | 54 hours ($n$ = 2) | 1.5 ± 0.71 (*not included in density analyses*) |
| | 60 hours ($n$ = 4) | 4.75 ± 0.50 |
| | 67 hours ($n$ = 4) | 63.0 ± 29.45 |

### 2.5 Cortical Mapping

Brain and spinal cord sections were examined using bright-field microscopy and polarized light illumination. To create cortical maps, a computer-based charting system was used to delineate section outlines, rabies-labeled neurons, gray-white matter boundaries, and other neuroanatomical features (i.e., corpus callosum, M1/S1 border, M1/M2 border, rhinal fissure) from every other section of the mouse brain as described previously (18). The nomenclature and boundaries for cortical areas were based on a standard atlas of the mouse brain (19). Custom laboratory software enabled the interactive aligning of charts of these individual sections. An “unfolding” line was then drawn along the top of cortical layer V and labeled neurons were projected to that line. The entire sequence of unfolded lines was then combined to generate high-resolution unfolded cortical maps as described previously (18). These maps, developed for both the TA and soleus muscles, display the location of labeled neurons in a two-dimensional representation of the cerebral cortex. Reconstructed cortical maps from experimental animals were aligned and scaled to a standardized atlas using common features such as the locations of the midline and the agranular-granular border between M1 and S1 as described previously in rats (20). Density-based analyses were performed to assess areas with peak labeling after peripheral muscle injections. To determine the density of labeling, we counted infected neurons in successive 100 µm by 100 µm bins.

### 2.6 Statistical Analyses

Statistical analyses were performed using GraphPad Prism 9 (GraphPad Software, San Diego, CA). Comparisons between any two groups of data were accomplished using the unpaired t test. Comparisons involving multiple groups were accomplished using two-way analysis of variance (ANOVA) followed by Bonferroni post-hoc analysis. All data are presented as mean ± SD. A *p* value ≤ 0.05 was considered statistically significant.

## 3. Results

The N2c strain of rabies virus is transported trans-synaptically in the retrograde direction via motor efferent pathways following injection into peripheral muscles (21). We used this approach to identify the anatomical nodes within the brain and spinal cord with the most direct synaptic connections to the motor neurons responsible for innervating hindlimb muscles in the mouse. Following unilateral virus injections into either the tibialis anterior (TA) or soleus muscle, multiple stages of viral replication and transneuronal transport resulted in the sequential infection of synaptically interconnected neurons (**Figure 1**). Through careful adjustment of survival time (24-93 hours), rabies-infected cells were first observed in spinal motor neurons and, subsequently, in spinal interneurons and brainstem nuclei prior to transport to cortical layer V and then cortical layers II/III and VI. Animals did not display symptoms during the extended survival periods used in these experiments, consistent with previous reports using rabies virus in the rodent (20).

**Figure 1.**
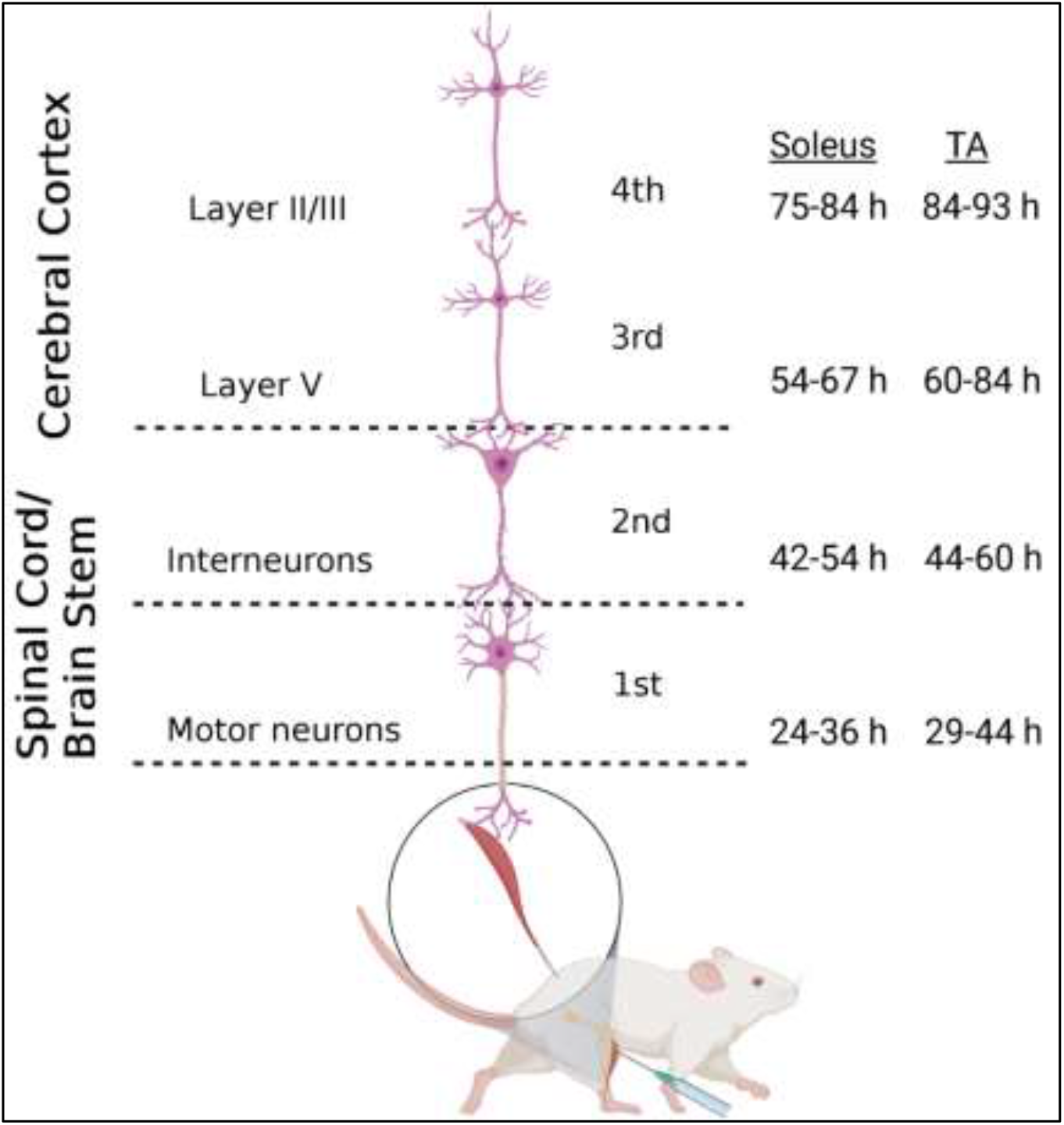
Retrograde transneuronal transport of rabies virus through the neural circuits that innervate muscles of the hindlimb. Following intramuscular injection, rabies virus is transported through a chain of synaptically-connected neurons. Once injected into a peripheral muscle (i.e., tibialis anterior or soleus), retrograde transport of the virus results in infection of motor neurons in the ventral horn of the spinal cord (1^st^ order) then interneurons in the intermediate zone of the spinal cord and/or in the brainstem (2^nd^ order) and then neurons in layer V of the cerebral cortex (3^rd^ order) in a time-dependent manner. From layer V of the cerebral cortex, layer II/III neurons of the cerebral cortex, as well as other areas projecting to layer V, are infected (4^th^ order). The observed time course of this retrograde labeling at each location is summarized for soleus and tibialis anterior (TA). Each stage of transport in the figure is numbered as described in the text. Figure created with BioRender.com.

### 3.1 Cortical representations of hindlimb muscles

The first infected cortical neurons observed were located in layer V (**Supplemental Figure 1**), a major output layer of the cerebral cortex. By increasing the survival time, an additional stage of retrograde transneuronal transport allowed for the infection of fourth-order neurons located in other cortical layers, specifically layers II/III and layer VI. As noted in prior studies in the rodent (20), the presence of rabies-positive neurons in layer V and the absence of rabies-positive cells in layers II/III is “an unequivocal marker that transport is restricted to those cortical neurons that are most directly connected to the muscle of interest.”

For animals with labeling restricted to third-order, infected cortical neurons were located in layer V only and predominantly in the hemisphere contralateral to the injection site (100%, 81%, and 74% for TA at 63-69 hours, 75-84 hours, and 84 hours, respectively; 95% and 86% for soleus at 60 hours and 67 hours, respectively). Following transport from TA, the cortical areas with infected neurons 63-69 hours post-injection were the primary motor cortex (M1, 64%), the rostromedial motor area (M2, 12%), and the primary somatosensory cortex (S1, 24%). This distribution expanded to include additional cortical areas 75-84 hours post-injection [M1 (45%), M2 (12%), S1 (41%), S2 (1%), and Cg1 (1%)] and 84 hours post-injection [M1 (52%), M2 (16%), S1 (30%), S2 (1%), and Cg1 (1%)] (**Figure 2** and **Supplemental Figure 2**). Similarly, following transport from soleus muscle, cortical layer V labeling was first observed at 60 hours with peak labeling observed at 67 hours. At 60 hours, the cortical areas with infected neurons were M1 (47%), M2 (16%), and S1 (37%) with similar cortical areas labeled at 67 hours post-injection [M1 (47%), M2 (9%), and S1 (44%)] (**Figure 3** and **Supplemental Figure 3**). These motor and non-motor areas represent the most direct cortical influences on both TA and soleus.

**Figure 2.**
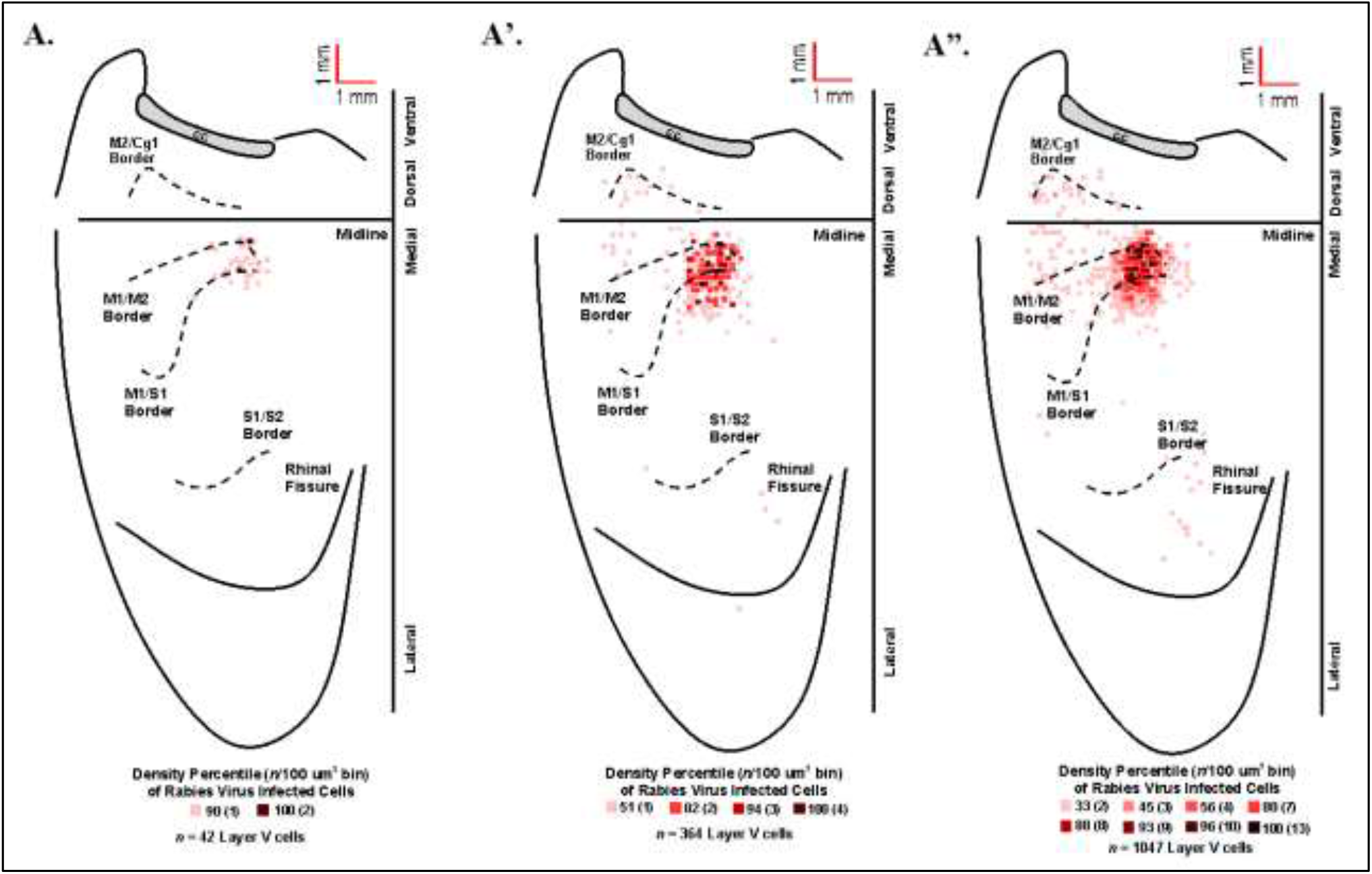
Origin of cortical projections to tibialis anterior (TA) motor neurons. Density-based analysis of neuronal labeling in cortical layer V of the contralateral hemisphere after retrograde transneuronal transport of RV from the right TA muscle [**A**: 63-69 hours (*n* = 5); **A’**: 75-84 hours (*n* = 4); **A**”: 84 hours (*n* = 2)]. Each square indicates the density of the infected neurons in bins (100 µm x 100 µm) throughout the cerebral cortex. Different colored bins represent percentiles of labeled cells; darker colors are higher percentiles. The numbers in parenthesis indicate the number of neurons in each bin. The medial wall of the contralateral hemisphere has been reflected upward and joined to the lateral surface at the midline. Each density map is a composite of multiple experiments that were overlapped on the atlas template. Cytoarchitectonic borders are delineated including the border between granular (S1) and agranular (M1) cortex in the region of the forelimb and hindlimb representations. Midline, midline of the hemisphere; M1, primary motor cortex; M2, secondary motor cortex (rostromedial motor field); S1, primary somatosensory cortex; S2, secondary somatosensory cortex; Cg1, cingulate cortex; CC, corpus callosum.

**Figure 3.**
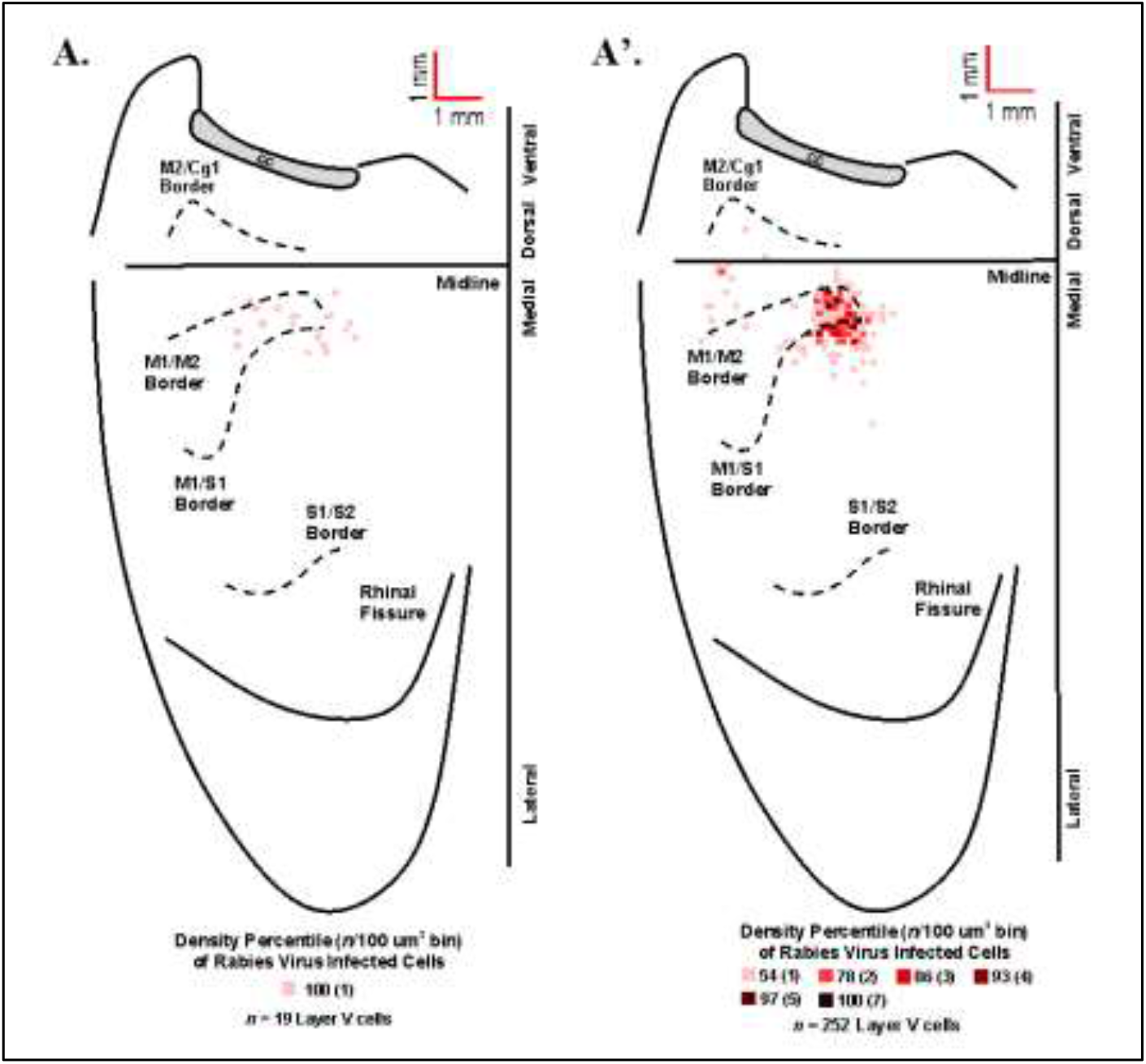
Origin of cortical projections to soleus motor neurons. Density-based analysis of neuronal labeling in cortical layer V of the contralateral hemisphere after retrograde transneuronal transport of RV from the right soleus muscle [**A**: 60 hours (*n* = 4); **A’**: 67 hours (*n* = 4)]. Each square indicates the density of the infected neurons in bins (100 µm x 100 µm) throughout the cerebral cortex. Conventions and abbreviations as in Figure 2.

Rabies-positive neurons were observed in additional cortical layers when survival times were extended to allow for fourth-order labeling. These infected neurons in the supragranular cortical layers were localized predominantly in M1, M2, and S1 in regions comparable to those observed for third-order neurons in layer V at shorter survival times (**Figure 4**). Taken together, these observations indicate that M1, M2, and S1 are the origin of the most direct cortical control of these two hindlimb muscles and that this cortical control is mediated by interneurons located in the brainstem and spinal cord.

**Figure 4.**
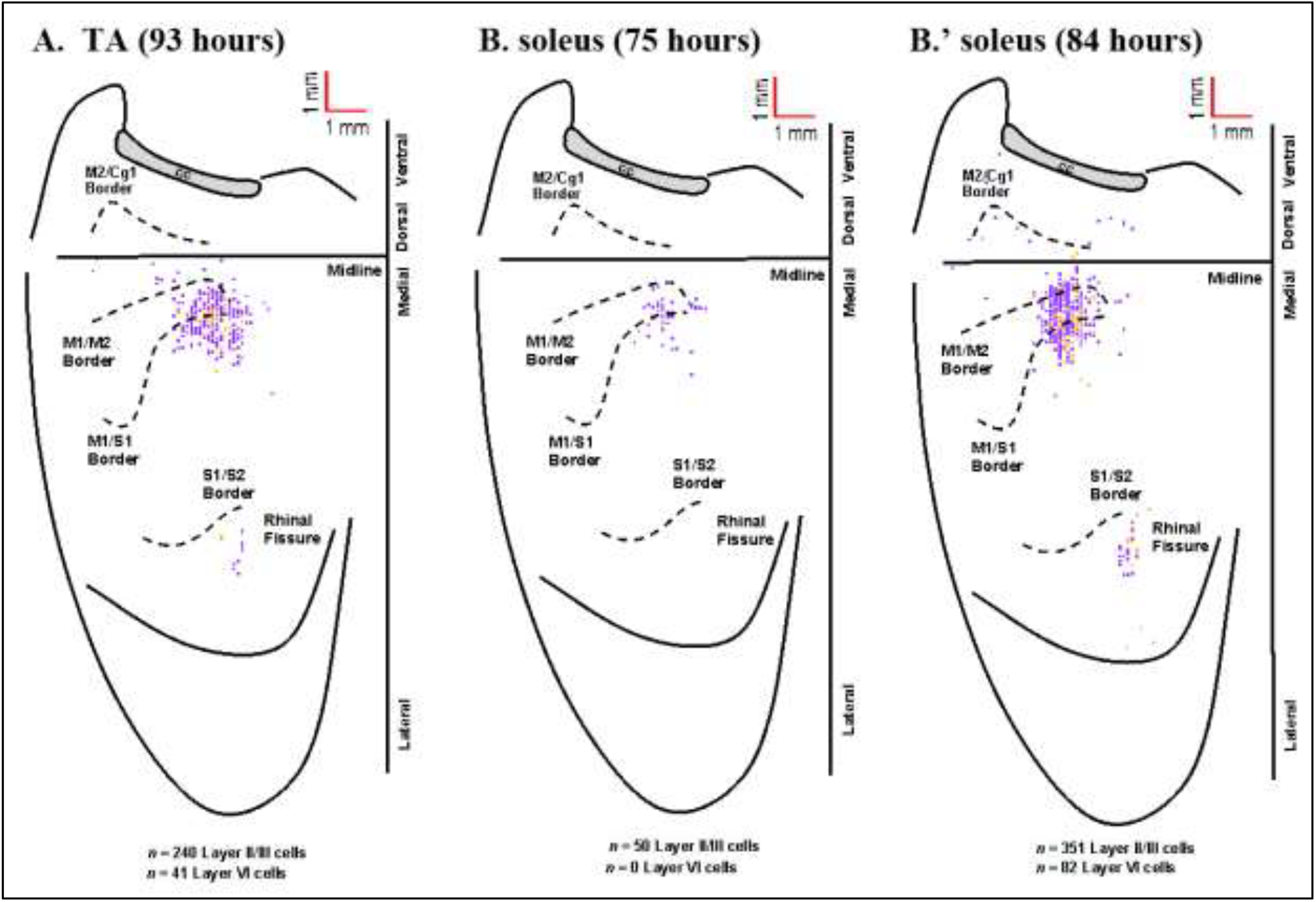
Cortical maps from animals with fourth-order labeling. Fourth-order cortical neurons are located most densely in the limb representations of M1, S1, and M2. **A**: Following RV injection into TA, infected neurons were observed in supragranular layers (layer IV and above, purple) and in layer VI (gold) at the 93-hour survival time. **B**: Following RV injection into soleus, infected neurons were observed in supragranular layers (layer IV and above, purple) at the 75-hour survival time. **B’**: At the 84-hour survival time, infected neurons were observed in supragranular layers (layer IV and above, purple) and in layer VI (gold) following RV injection into soleus. All maps are from representative animals. Each square represents a single labeled neuron. Conventions and abbreviations as in Figure 2.

### 3.2 Cortical representations of fast-versus slow-twitch muscles

Fiber-type impacts a number of aspects of muscle physiology including energy usage, fatiguability, and recruitment order. To determine if differences in innervation patterns between fast- and slow-twitch muscles exist, we compared the anatomical nodes and neuronal populations labeled following rabies virus injection and found that fiber-types also impacts viral uptake. First, we observed spinal motor neuron labeling at shorter survival times after RV injections into the slow-twitch soleus muscle when compared to the primarily fast-twitch TA muscle. Second, we describe differences in the time course of cortical labeling following soleus and TA injections that appear to reflect their differential muscle fiber composition.

Overall, the anatomical location of the cortical neurons innervating the soleus and TA were consistent across the two muscles. In addition, the cortical representations in M1 showed significant overlap. However, the labeling after TA injections was somewhat more extensive than after soleus injections, in particular in S1. This is best observed when the cortical labeling associated with each muscle is overlaid (**Figure 5**).

**Figure 5.**
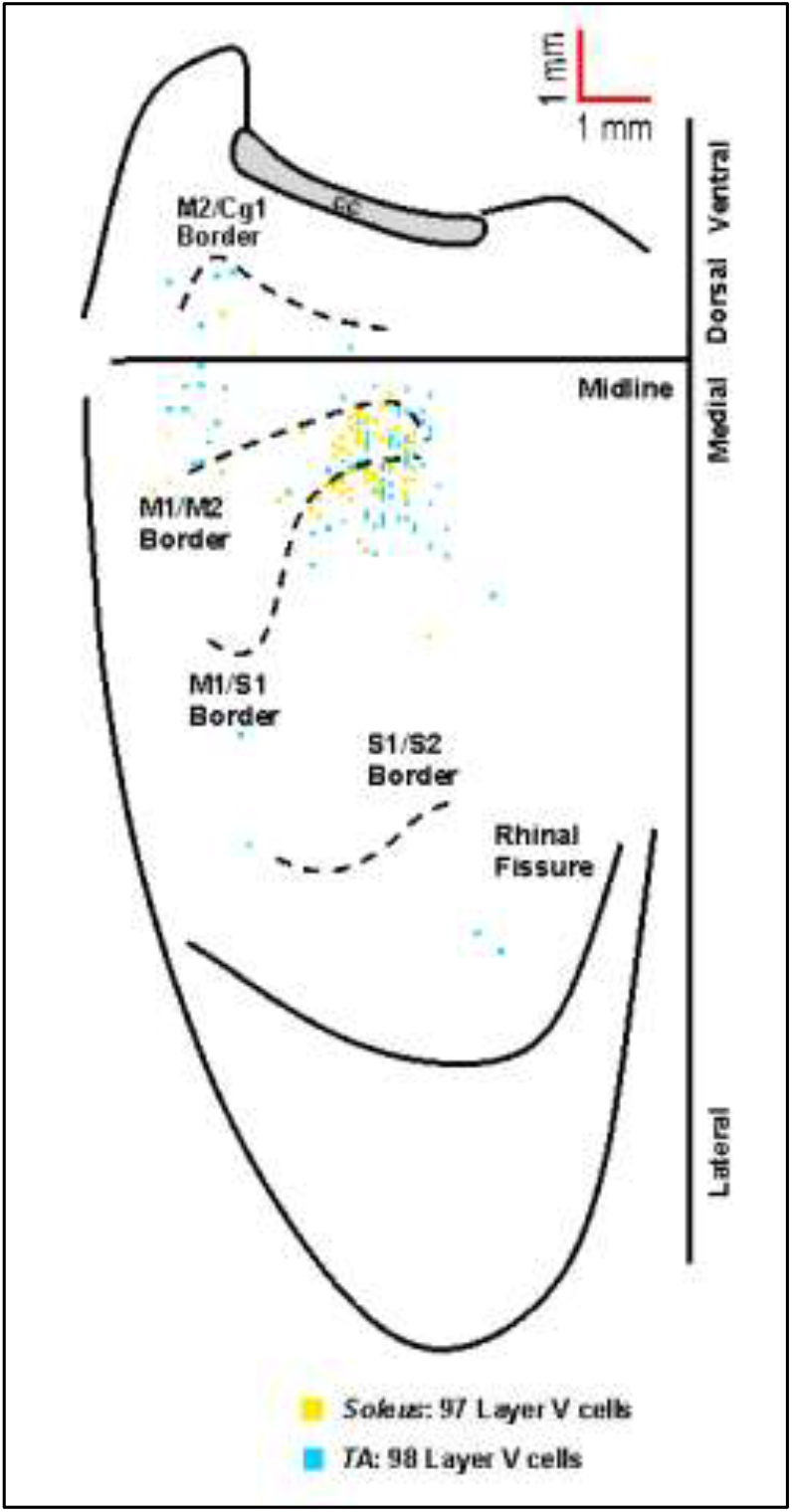
Overlay of the cortical representations of TA and soleus muscles. Representative cortical maps from animals with third-order neuronal labeling following RV injection into TA (blue) or soleus (yellow) show the origin of projections to each hindlimb muscle. The medial wall of the contralateral hemisphere has been reflected upward and joined to the lateral surface at the midline. Each square represents a single labeled neuron. Cortical projections originate from overlapping regions of M1, M2, and S1 for TA and soleus. Conventions and abbreviations as in Figure 2.

### 3.3 Anatomical nodes and timing of retrograde transneuronal transport

Spinal motor neuron labeling was assessed as the first relevant anatomical node and was used to establish the lower range of survival times necessary for transport (**Supplemental Figure 1**). Labeling of these cells is associated with the retrograde transport of rabies virus from injected muscles via the neuromuscular junction to the motor neurons directly responsible for their control. First-order spinal motor neurons were observed in lumbar segments (approximately L4-L5 for TA and approximately L3-L4 for soleus) with labeling first noted at 24 hours for soleus and at 29 hours for TA.

The transition to second-order labeling was considered to be when the labeling of spinal cord interneurons and/or brainstem neurons was observed and indicates the retrograde transneuronal transport of rabies virus from motor neurons to neuronal cells with direct synaptic communication with these cells. This labeling was apparent at longer survival times (44-60 hours for TA and 42-54 hours for soleus) and included neurons in both the ipsilateral and contralateral spinal cord, as well as the area surrounding the central canal. Brainstem second-order premotor neurons were observed predominantly in the magnocellular reticular nucleus (Mc), pontine reticular nucleus (Pn), gigantocellular reticular nucleus (Gi), spinal vestibular nucleus (SpVe) and vestibular nucleus (Ve) following hindlimb muscle injections. There were no apparent differences in the extent of labeling at these sites following TA or soleus injections.

Third-order cortical labeling was subsequently observed at survival times of 60-84 hours for TA and 54-67 hours for soleus. The time course of this transport process was established for each hindlimb muscle. Initial comparisons indicated that first-order labeling was observed at shorter survival times following viral injections for the soleus muscle compared to TA (soleus 30.55 ± 4.99 hours, TA 37.46 ± 5.59 hours, p < 0.01; **Figure 6**, black and gray bars). Similar differences between TA and S were also observed at higher orders of transport—second order: soleus 46.80 ± 4.73 hours, TA 55.06 ± 6.07 hours, p < 0.001; third order: soleus 61.60 ± 5.19 hours, TA 71.58 ± 9.07 hours, p < 0.01; and fourth order: soleus 79.50 ± 4.81 hours, TA 91.2 ± 4.02 hours, p < 0.001 (**Figure 6**, black and gray bars). To determine if these differences were due to a differential rate of virus uptake from the predominantly (90% type II) fast-twitch TA versus the slow-twitch (50% type I) soleus, we normalized the higher order survival times by subtracting the mean survival time for first order labeling (37.46 hours for TA; 30.55 hours for soleus) from the survival times associated with second, third, and fourth order labeling. After normalization, the rates of retrograde transneuronal transport from TA or soleus were not significantly different (second order: p = 0.565; third order: p = 0.362; fourth order: p = 0.0094; **Figure 6**, green and light blue bars). This analysis indicates that once the virus is transported retrogradely from the muscle to the motor neurons, further retrograde transneuronal transport occurs at the same rate regardless of the muscle of origin.

**Figure 6.**
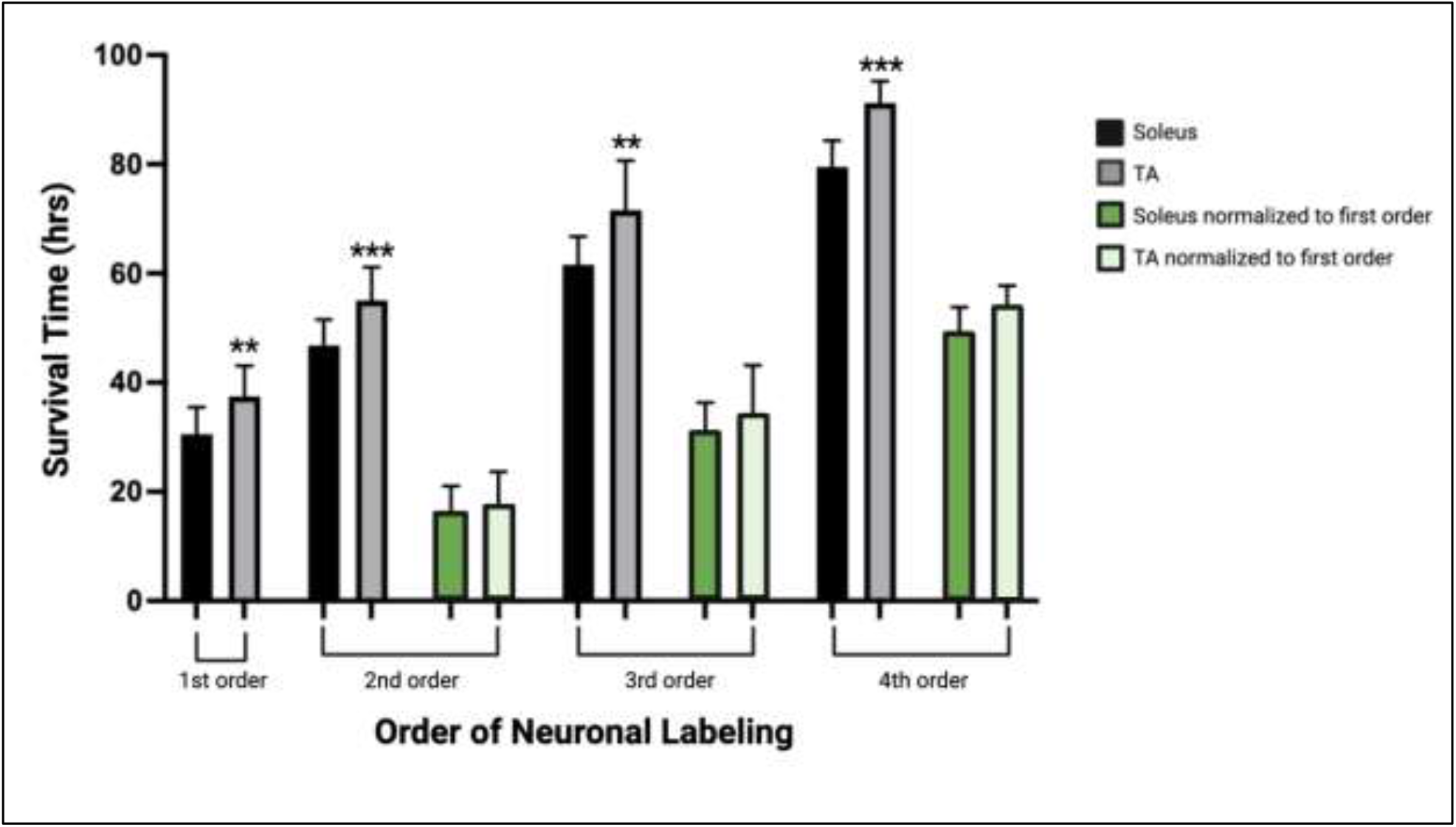
The time course of transport to locations throughout the motor system (orders of labeling as shown in Figure 1) following viral injection. Mean survival times (hours) and standard deviations associated with first, second, third, and fourth order labeling are provided (black bars, soleus; gray bars, TA). To delineate viral uptake at the neuromuscular junction from transneuronal transport, the timing of second, third, and fourth order labeling were normalized to the mean survival time associated with first order labeling (30.55 hours for soleus and 37.46 hours for TA; green bars, soleus; light blue bars, TA). **p < 0.01; ***p < 0.001.

### 3.4 Efficiency of spinal motor neuron and cortical layer V transduction

Rabies virus is a rhabdovirus with a marked neurotropism for the muscular form of the nicotinic acetylcholine receptor (22). This neurotropism facilitates viral entry into cells at the neuromuscular junction. To assess the efficiency of transduction after injection, we evaluated the number of spinal motor neurons labeled prior to interneuron labeling (**Table 1**). For TA, this first-order labeling occurred at survival times ranging from 29-44 hours. The estimated percentage of the TA spinal motor neuron pool labeled was 3.3% at 29 hours, 9.8% at 36 hours, 12.7% at 40 hours, and 7.0% at 44 hours. The amount of labeling reached a maximum level between 36 and 40 hours, with no increase at the longer survival time. First-order labeling following soleus muscle injections was observed at the 24-hour survival time, with second order interneuron labeling observed as early as 42 hours after viral injections. The estimated percentage of the spinal motor neuron pool for soleus labeled was 39.0% at 29 hours, 56.0% at 30 hours, and 54.8% at 36 hours. For soleus, the amount of labeling reached a maximum by 30 hours.

Following viral infection of motor neurons, rabies virus was transported trans-synaptically in the retrograde direction to interneurons (2^nd^ order transport) and then, after another order of transneuronal transport, to cortical neurons in layer V (3^rd^ order transport). Cortical layer V neurons are the cortical neurons with the most direct cortical projections to each muscle’s motor neurons. The earliest layer V labeling was observed at 60 hours after TA injections and at 54 hours following soleus injections. The number of layer V neurons labeled in those animals without 4^th^ order labeling (i.e., labeling in layers II/III or VI) were counted (**Table 2**). For both muscles, layer V labeling continued to increase at longer survival times until the survival time at which labeling in cortical layers II/III and VI (4^th^ order) was noted.

## 4. Discussion

In this study, we used retrograde transneuronal transport of rabies virus to identify and compare the cortical areas that control fast- and slow-twitch hindlimb muscles surrounding the ankle joint of the mouse and reported the density-based distribution of these cortical connections. Viral uptake from the slow-twitch soleus muscle injection and subsequent labeling of spinal motor neurons occurred at a faster rate than viral uptake and spinal motor neuron labeling from the fast-twitch TA muscle injection. However, we established that the time course for further retrograde transneuronal transport of rabies virus from spinal motor neurons to neurons in the cerebral cortex does not differ between the two muscles. Our data provides a foundation for future studies aimed at understanding disease-specific changes that occur in particular cellular populations within these neural networks and will specifically help to identify pathological alterations related to retrograde transneuronal transport.

Despite its central role in the study of disease, the mouse motor system has yet to be fully characterized, particularly with respect to hindlimb muscles that are less involved in the completion of sophisticated tasks (4). In this study, the cortical areas associated with the control of hindlimb muscles were consistent with those noted in stimulation studies. In particular, the hindlimb was represented in the caudal portion of M1 and medial to S1, although overlap with the sensory cortex was also noted (23, 24). Moreover, this hindlimb representation was located posterior to bregma and extended caudally as noted by others (25, 26). Most of the labeled layer V neurons (3^rd^ order) were located in the hindlimb representations of M1, M2, and S1 with additional labeling in regions (S2) surrounding the rhinal fissure. Third-order labeling of layer V neurons following TA injection was first observed at 60 hours and increased at longer survival times, reaching a peak at 84 hours. When considered in the context of the staggered temporal infection reported previously for the TA motor neuron pool (27), it is likely that this graded increase in cortical labeling after TA injections reflects progressive virus uptake from gamma, small alpha, and then large alpha motor neuron populations of the TA muscle. Similar results were obtained following soleus muscle injections with early cortical labeling observed at 54 hours with peak density of layer V neurons at 67 hours. This pattern of labeling may correspond to progressive virus uptake from the gamma and then small alpha motor neurons of the soleus muscle. Further studies are needed to more fully explore the relationship between muscle fiber composition and virus uptake and transport.

The overlap of the cortical representations of the two hindlimb muscles (**Figure 5**) has several important implications. First, we observed a clear separation between the hindlimb representation and areas involved in forelimb control (23), consistent with previous findings in both rodents and non-human primates (3). In addition, our data indicate that cortical representations of single hindlimb muscles in mouse motor cortex are not unique, but rather exhibit considerable overlap and intermingling with other hindlimb muscles. This organization is in line with previous observations in the mouse forelimb (5) and with studies in the non-human primate using anatomical tracing to label single corticomotoneuronal (CM) cells (21) and stimulation to evoke muscle activity (28-34). Second, our results indicate that hindlimb muscles with opposing physiological actions (i.e., ankle flexion for TA versus ankle extension for soleus) occupy the same cortical space in the mouse, consistent with findings in non-human primates (35). Studies in non-human primates looking specifically at forelimb antagonist muscle representations have demonstrated that although these neurons occupy the same space in the cortex, they do not have overlapping branching patterns. In fact, Griffin *et al*. found that “…the multiple functions of a target muscle are represented by the activity of separate populations of CM cells” (36). Together with our findings, these results support the conclusion that corticospinal (CS) neurons differ not only on the specific muscles and muscle fiber types to which they project but also on their functional utilization. Future studies using dual-tracer approaches in the same animal and/or physiology techniques are needed to determine whether individual CS neurons map onto functionally distinct muscle types (e.g., flexors versus extensors) in the mouse motor cortex. Third, our results have important implications for pathological conditions like ALS and highlight that mechanisms beyond anatomical organization must be considered in the context of disease susceptibility. In particular, differential neuronal loss within the same cortical space suggests that circuitry-, cellular-, and molecular-level factors may play a role in disease susceptibility. While muscle fiber type impacts multiple aspects of muscle physiology including energy usage, fatiguability, recruitment order, and rabies virus uptake (as observed in the present study), additional mechanisms (perhaps within the spinal cord and/or based on inherent cellular differences) may also have an impact. Our ongoing studies seek to evaluate the relationship between these factors and disease susceptibility.

TA motor neurons were first labeled 29 hours following rabies virus injections into the muscle, and the number of motor neurons increased until 44 hours post-injection when second order labeling was first noted in spinal interneurons. These results are consistent with those obtained using pseudorabies virus (PRV)-Bartha, a DNA virus. With PRV-Bartha, the TA motor neuron pool was first labeled 24-28 hours following virus injection with increased labeling observed up to 72 hours (27). While the time course was similar, the number of motor neurons labeled with rabies virus was lower than that observed by others (16, 27, 37). The counts reported in our study may be decreased as a result of the limited amount of virus injected into each muscle to minimize its spread and/or the limited uptake of virus at sites distal to the neuromuscular junction. In contrast, rabies virus injections into the soleus muscle labeled the majority of motor neurons by 30 hours post-injection. Increasing the volume and number of injections into each muscle may be necessary to label a larger percentage of these cells. This will be explored in future studies such that further characterization of the size distribution and motor neuron subtypes can be accomplished.

In this study, we have carefully adjusted survival times to optimize transport to the cerebral cortex and did not pursue a precise definition of the specific connectivity (38) and microcircuit organization (39, 40) of spinal interneuron and brainstem populations. In addition, we noted variability in the order of labeling across animals as reported previously both in vitro and in vivo [reviewed in (13)]. However, we evaluated the time course of labeling in interneuron populations located both ipsilateral and contralateral to the injection site. As reported with PRV (27), ipsilateral interneurons were first observed dorsal, ventral, and medial to the motor neuron pools of both hindlimb muscles, particularly in laminae V-VIII and near the central canal. This ipsilateral labeling occurred 44 hours post-injection for TA with contralateral labeling in the spinal cord observed as early as 48 hours. Following soleus muscle injections, this ipsilateral interneuron labeling was observed as early as 42 hours, and contralateral interneuron labeling was consistently observed 48 hours after soleus injections. We hypothesize that these interneuron populations include Renshaw cells and Ia inhibitory neurons as both are known to have direct synaptic projections to motor neurons (41) and future analyses will include characterization of these subpopulations.

Brainstem labeling was observed prior to cortical labeling following viral injections into both hindlimb muscles. These observations are consistent with the work of others indicating that brainstem regions have monosynaptic connections to spinal motor neurons and serve as a relay station between the spinal cord and brain (4, 42). It also suggests the mouse has “last-order interneurons” (43) with motor cortex projections to spinal cord and brainstem regions as described in the rat (44-46). In other words, second-order neurons located in the spinal cord and brainstem mediate transport to third-order neurons located predominantly in M1, the region with the highest percentage of cortical cells labeled. Relatedly, the cortical labeling observed in layers II/III (i.e., fourth order labeling, **Figure 4**) was localized to regions associated with the labeling observed in layer V (i.e., third-order labeling, **Figures 2 and 3**) at the shortest survival times. As noted in the rat motor cortex (47), these hindlimb motor representations overlap with the corresponding sensory representations in what has been termed an “overlap zone,” characterized by granule cells in layer IV and pyramidal cells in layer V (23).

Overall, our results provide a foundation for ongoing work aimed at better understanding the impact of disease, particularly for conditions with pathological alterations of projections to and from the motor cortex (48) or axonal transport deficits. The ability to evaluate reorganization in specific motor networks may also reveal aspects related to differential neuronal vulnerability and the impact of cell death at particular nodes in the network. For example, there is evidence that alpha and gamma motor neurons are differentially affected in diseases of the motor system (49) and that the loss of specific spinal interneuron populations can lead to motor dysfunction (50). The ability to further investigate the associated circuitry in murine models may provide insights into these differences with potential avenues and targets for therapeutic interventions.

## Acknowledgments

The authors wish to thank Jean Alban Rathelot, PhD, Katherine Rohde, Riya Tipnis, Amina Chtourou, and Andrew Becker for technical assistance at earlier stages of this work; David Levinthal, MD, PhD for thoughtful discussions related to cortical mapping; Mike Page for software assistance; Richard Dum, PhD and Andreea Bostan, PhD for multiple thorough reviews and constructive comments; Darcy Griffin, PhD for expert talks on muscle representations in the cortex of non-human primates; and Peter Strick, PhD for critical review and comments during manuscript preparation. In addition, the authors wish to acknowledge the Center for Neuroanatomy with Neurotropic Viruses (CNNV; Peter Strick, PhD, Director) and the Center for Biologic Imaging (CBI; Simon Watkins, PhD, Director). Finally, the authors are grateful to J. Cole for inspiring the completion of this work done in memory of Ronald Maurer.

## Author Contributions Statement

CK was responsible for the conception of the study and for the experimental design. CK, LM, MB, and TS contributed to the acquisition, analysis, and interpretation of the data including statistical analyses and figure preparation. CK wrote the first draft of the manuscript. MB, LM, and TS reviewed sections of the manuscript. All authors contributed to manuscript revision and have read and approved of the submitted version.

## Funding

Funding for this work was provided by the Muscular Dystrophy Association and the American Association of Neuromuscular and Electrodiagnostic Medicine (CK; Development Grant Award Number: 416035) and The ALS Association (CK; Investigator-Initiated Starter Award Number: 20-IIP-512). Additional financial support was provided by the Center for Neuroanatomy with Neurotropic Viruses (CNNV; P40 OD010996; PI: Strick) and from the University of Pittsburgh Brain Institute.

## Notes

Conflict of Interest Statement: The authors declare that the research was conducted in the absence of any commercial or financial relationships that could be construed as a potential conflict of interest.

### Competing Interest Statement

The authors have declared no competing interest.

## References

1. Lemon RN. Descending pathways in motor control. Annu Rev Neurosci. 2008;31:195–218. Epub 2008/06/19. doi: 10.1146/annurev.neuro.31.060407.125547. PubMed PMID: 18558853.

2. Rathelot JA, Strick PL. Subdivisions of primary motor cortex based on cortico-motoneuronal cells. Proc Natl Acad Sci U S A. 2009;106(3):918–23. Epub 2009/01/14. doi: 10.1073/pnas.0808362106. PubMed PMID: 19139417; PMCID: PMC2621250.

3. Strick PL, Dum RP, Rathelot JA. The Cortical Motor Areas and the Emergence of Motor Skills: A Neuroanatomical Perspective. Annu Rev Neurosci. 2021;44:425–47. Epub 2021/04/18. doi: 10.1146/annurev-neuro-070918-050216. PubMed PMID: 33863253.

4. Esposito MS, Capelli P, Arber S. Brainstem nucleus MdV mediates skilled forelimb motor tasks. Nature. 2014;508(7496):351–6. Epub 2014/02/04. doi: 10.1038/nature13023. PubMed PMID: 24487621.

5. Ueno M, Nakamura Y, Li J, Gu Z, Niehaus J, Maezawa M, Crone SA, Goulding M, Baccei ML, Yoshida Y. Corticospinal Circuits from the Sensory and Motor Cortices Differentially Regulate Skilled Movements through Distinct Spinal Interneurons. Cell Rep. 2018;23(5):1286–300 e7. Epub 2018/05/03. doi: 10.1016/j.celrep.2018.03.137. PubMed PMID: 29719245; PMCID: PMC6608728.

6. Henneman E, Somjen G, Carpenter DO. Functional Significance of Cell Size in Spinal Motoneurons. J Neurophysiol. 1965;28:560–80. Epub 1965/05/01. doi: 10.1152/jn.1965.28.3.560. PubMed PMID: 14328454.

7. Ciciliot S, Rossi AC, Dyar KA, Blaauw B, Schiaffino S. Muscle type and fiber type specificity in muscle wasting. Int J Biochem Cell Biol. 2013;45(10):2191–9. Epub 2013/05/25. doi: 10.1016/j.biocel.2013.05.016. PubMed PMID: 23702032.

8. Frey D, Schneider C, Xu L, Borg J, Spooren W, Caroni P. Early and selective loss of neuromuscular synapse subtypes with low sprouting competence in motoneuron diseases. J Neurosci. 2000;20(7):2534–42. Epub 2000/03/24. PubMed PMID: 10729333; PMCID: PMC6772256.

9. Pun S, Santos AF, Saxena S, Xu L, Caroni P. Selective vulnerability and pruning of phasic motoneuron axons in motoneuron disease alleviated by CNTF. Nat Neurosci. 2006;9(3):408–19. Epub 2006/02/14. doi: 10.1038/nn1653. PubMed PMID: 16474388.

10. Hegedus J, Putman CT, Gordon T. Time course of preferential motor unit loss in the SOD1 G93A mouse model of amyotrophic lateral sclerosis. Neurobiol Dis. 2007;28(2):154–64. Epub 2007/09/04. doi: 10.1016/j.nbd.2007.07.003. PubMed PMID: 17766128.

11. Ugolini G. Advances in viral transneuronal tracing. J Neurosci Methods. 2010;194(1):2–20. Epub 2009/12/17. doi: 10.1016/j.jneumeth.2009.12.001. PubMed PMID: 20004688.

12. Kelly RM, Strick PL. Cerebellar loops with motor cortex and prefrontal cortex of a nonhuman primate. J Neurosci. 2003;23(23):8432–44. Epub 2003/09/12. PubMed PMID: 12968006; PMCID: PMC6740694.

13. Kelly RM, Strick PL. Rabies as a transneuronal tracer of circuits in the central nervous system. J Neurosci Methods. 2000;103(1):63–71. Epub 2000/11/14. doi: 10.1016/s0165-0270(00)00296-x. PubMed PMID: 11074096.

14. Charles JP, Cappellari O, Spence AJ, Hutchinson JR, Wells DJ. Musculoskeletal Geometry, Muscle Architecture and Functional Specialisations of the Mouse Hindlimb. PLoS One. 2016;11(4):e0147669. Epub 2016/04/27. doi: 10.1371/journal.pone.0147669. PubMed PMID: 27115354; PMCID: PMC4846001.

15. Hoshi E, Tremblay L, Feger J, Carras PL, Strick PL. The cerebellum communicates with the basal ganglia. Nat Neurosci. 2005;8(11):1491–3. Epub 2005/10/06. doi: 10.1038/nn1544. PubMed PMID: 16205719.

16. Tissenbaum HA, Parry DJ. The effect of partial denervation of tibialis anterior (TA) muscle on the number and sizes of motorneurons in TA motornucleus of normal and dystrophic (C57BL dy2j/dy2j) mice. Can J Physiol Pharmacol. 1991;69(11):1769–73. Epub 1991/11/01. doi: 10.1139/y91-261. PubMed PMID: 1804521.

17. Ishihara A, Ohira Y, Tanaka M, Nishikawa W, Ishioka N, Higashibata A, Izumi R, Shimazu T, Ibata Y. Cell body size and succinate dehydrogenase activity of spinal motoneurons innervating the soleus muscle in mice, rats, and cats. Neurochem Res. 2001;26(12):1301–4. Epub 2002/03/12. doi: 10.1023/a:1014245417017. PubMed PMID: 11885781.

18. Dum RP, Strick PL. The origin of corticospinal projections from the premotor areas in the frontal lobe. J Neurosci. 1991;11(3):667–89. Epub 1991/03/01. PubMed PMID: 1705965; PMCID: PMC6575356.

19. Paxinos Ga f, K. The mouse brain in stereotaxic coordinates. Fifth ed 2019 April 6, 2019.

20. Levinthal DJ, Strick PL. The motor cortex communicates with the kidney. J Neurosci. 2012;32(19):6726–31. Epub 2012/05/11. doi: 10.1523/JNEUROSCI.0406-12.2012. PubMed PMID: 22573695; PMCID: PMC3363289.

21. Rathelot JA, Strick PL. Muscle representation in the macaque motor cortex: an anatomical perspective. Proc Natl Acad Sci U S A. 2006;103(21):8257–62. Epub 20060515. doi: 10.1073/pnas.0602933103. PubMed PMID: 16702556; PMCID: PMC1461407.

22. Lentz TL, Burrage TG, Smith AL, Tignor GH. The acetylcholine receptor as a cellular receptor for rabies virus. Yale J Biol Med. 1983;56(4):315–22. Epub 1983/07/01. PubMed PMID: 6367238; PMCID: PMC2589619.

23. Tennant KA, Adkins DL, Donlan NA, Asay AL, Thomas N, Kleim JA, Jones TA. The organization of the forelimb representation of the C57BL/6 mouse motor cortex as defined by intracortical microstimulation and cytoarchitecture. Cereb Cortex. 2011;21(4):865–76. Epub 2010/08/27. doi: 10.1093/cercor/bhq159. PubMed PMID: 20739477; PMCID: PMC3059888.

24. Ayling OG, Harrison TC, Boyd JD, Goroshkov A, Murphy TH. Automated light-based mapping of motor cortex by photoactivation of channelrhodopsin-2 transgenic mice. Nat Methods. 2009;6(3):219–24. Epub 2009/02/17. doi: 10.1038/nmeth.1303. PubMed PMID: 19219033.

25. Pronichev IV, Lenkov DN. Functional mapping of the motor cortex of the white mouse by a microstimulation method. Neurosci Behav Physiol. 1998;28(1):80–5. Epub 1998/03/26. doi: 10.1007/BF02461916. PubMed PMID: 9513982.

26. Li CX, Waters RS. Organization of the mouse motor cortex studied by retrograde tracing and intracortical microstimulation (ICMS) mapping. Can J Neurol Sci. 1991;18(1):28–38. Epub 1991/02/01. doi: 10.1017/s0317167100031267. PubMed PMID: 2036613.

27. Jovanovic K, Pastor AM, O’Donovan MJ. The use of PRV-Bartha to define premotor inputs to lumbar motoneurons in the neonatal spinal cord of the mouse. PLoS One. 2010;5(7):e11743. Epub 2010/07/30. doi: 10.1371/journal.pone.0011743. PubMed PMID: 20668534; PMCID: PMC2909228.

28. Hudson HM, Park MC, Belhaj-Saif A, Cheney PD. Representation of individual forelimb muscles in primary motor cortex. J Neurophysiol. 2017;118(1):47–63. Epub 2017/03/31. doi: 10.1152/jn.01070.2015. PubMed PMID: 28356482; PMCID: PMC5494364.

29. Gould HJ, 3rd, Cusick CG, Pons TP, Kaas JH. The relationship of corpus callosum connections to electrical stimulation maps of motor, supplementary motor, and the frontal eye fields in owl monkeys. J Comp Neurol. 1986;247(3):297–325. Epub 1986/05/15. doi: 10.1002/cne.902470303. PubMed PMID: 3722441.

30. Waters RS, Samulack DD, Dykes RW, McKinley PA. Topographic organization of baboon primary motor cortex: face, hand, forelimb, and shoulder representation. Somatosens Mot Res. 1990;7(4):485–514. Epub 1990/01/01. doi: 10.3109/08990229009144721. PubMed PMID: 2291379.

31. Donoghue JP, Leibovic S, Sanes JN. Organization of the forelimb area in squirrel monkey motor cortex: representation of digit, wrist, and elbow muscles. Exp Brain Res. 1992;89(1):1–19. Epub 1992/01/01. doi: 10.1007/BF00228996. PubMed PMID: 1601087.

32. Schieber MH. Constraints on somatotopic organization in the primary motor cortex. J Neurophysiol. 2001;86(5):2125–43. Epub 2001/11/08. doi: 10.1152/jn.2001.86.5.2125. PubMed PMID: 11698506.

33. Graziano MS, Taylor CS, Moore T. Complex movements evoked by microstimulation of precentral cortex. Neuron. 2002;34(5):841–51. Epub 2002/06/14. doi: 10.1016/s0896-6273(02)00698-0. PubMed PMID: 12062029.

34. Graziano MS, Aflalo TN, Cooke DF. Arm movements evoked by electrical stimulation in the motor cortex of monkeys. J Neurophysiol. 2005;94(6):4209–23. Epub 2005/08/27. doi: 10.1152/jn.01303.2004. PubMed PMID: 16120657.

35. Hudson HM, Griffin DM, Belhaj-Saif A, Cheney PD. Cortical output to fast and slow muscles of the ankle in the rhesus macaque. Front Neural Circuits. 2013;7:33. Epub 2013/03/06. doi: 10.3389/fncir.2013.00033. PubMed PMID: 23459919; PMCID: PMC3585439.

36. Griffin DM, Hoffman DS, Strick PL. Corticomotoneuronal cells are “functionally tuned”. Science. 2015;350(6261):667–70. Epub 2015/11/07. doi: 10.1126/science.aaa8035. PubMed PMID: 26542568; PMCID: PMC4829105.

37. Dekkers J, Bayley P, Dick JR, Schwaller B, Berchtold MW, Greensmith L. Over-expression of parvalbumin in transgenic mice rescues motoneurons from injury-induced cell death. Neuroscience. 2004;123(2):459–66. Epub 2003/12/31. doi: 10.1016/j.neuroscience.2003.07.013. PubMed PMID: 14698753.

38. Goulding M, Bourane S, Garcia-Campmany L, Dalet A, Koch S. Inhibition downunder: an update from the spinal cord. Curr Opin Neurobiol. 2014;26:161–6. Epub 2014/04/20. doi: 10.1016/j.conb.2014.03.006. PubMed PMID: 24743058; PMCID: PMC4059017.

39. Ziskind-Conhaim L, Hochman S. Diversity of molecularly defined spinal interneurons engaged in mammalian locomotor pattern generation. J Neurophysiol. 2017;118(6):2956–74. Epub 2017/09/01. doi: 10.1152/jn.00322.2017. PubMed PMID: 28855288; PMCID: PMC5712661.

40. Bikoff JB, Gabitto MI, Rivard AF, Drobac E, Machado TA, Miri A, Brenner-Morton S, Famojure E, Diaz C, Alvarez FJ, Mentis GZ, Jessell TM. Spinal Inhibitory Interneuron Diversity Delineates Variant Motor Microcircuits. Cell. 2016;165(1):207–19. Epub 2016/03/08. doi: 10.1016/j.cell.2016.01.027. PubMed PMID: 26949184; PMCID: PMC4808435.

41. Renshaw B. Central effects of centripetal impulses in axons of spinal ventral roots. J Neurophysiol. 1946;9:191–204. Epub 1946/05/01. doi: 10.1152/jn.1946.9.3.191. PubMed PMID: 21028162.

42. Pivetta C, Esposito MS, Sigrist M, Arber S. Motor-circuit communication matrix from spinal cord to brainstem neurons revealed by developmental origin. Cell. 2014;156(3):537–48. Epub 2014/02/04. doi: 10.1016/j.cell.2013.12.014. PubMed PMID: 24485459.

43. Brownstone RM, Bui TV. Spinal interneurons providing input to the final common path during locomotion. Prog Brain Res. 2010;187:81–95. Epub 2010/11/30. doi: 10.1016/B978-0-444-53613-6.00006-X. PubMed PMID: 21111202; PMCID: PMC3150186.

44. Ba-M’Hamed S, Roy JC, Bennis M, Poulain P, Sequeira H. Corticospinal collaterals to medullary cardiovascular nuclei in the rat: an anterograde and a retrograde double-labeling study. J Hirnforsch. 1996;37(3):367–75. Epub 1996/01/01. PubMed PMID: 8872559.

45. Miller MW. The origin of corticospinal projection neurons in rat. Exp Brain Res. 1987;67(2):339–51. Epub 1987/01/01. doi: 10.1007/BF00248554. PubMed PMID: 3622693.

46. Gabbott PL, Warner TA, Jays PR, Salway P, Busby SJ. Prefrontal cortex in the rat: projections to subcortical autonomic, motor, and limbic centers. J Comp Neurol. 2005;492(2):145–77. Epub 2005/10/01. doi: 10.1002/cne.20738. PubMed PMID: 16196030.

47. Donoghue JP, Wise SP. The motor cortex of the rat: cytoarchitecture and microstimulation mapping. J Comp Neurol. 1982;212(1):76–88. Epub 1982/11/20. doi: 10.1002/cne.902120106. PubMed PMID: 6294151.

48. Commisso B, Ding L, Varadi K, Gorges M, Bayer D, Boeckers TM, Ludolph AC, Kassubek J, Muller OJ, Roselli F. Stage-dependent remodeling of projections to motor cortex in ALS mouse model revealed by a new variant retrograde-AAV9. Elife. 2018;7. Epub 2018/08/24. doi: 10.7554/eLife.36892. PubMed PMID: 30136928; PMCID: PMC6125125.

49. Lalancette-Hebert M, Sharma A, Lyashchenko AK, Shneider NA. Gamma motor neurons survive and exacerbate alpha motor neuron degeneration in ALS. Proc Natl Acad Sci U S A. 2016;113(51):E8316–E25. Epub 2016/12/09. doi: 10.1073/pnas.1605210113. PubMed PMID: 27930290; PMCID: PMC5187676.

50. Salamatina A, Yang JH, Brenner-Morton S, Bikoff JB, Fang L, Kintner CR, Jessell TM, Sweeney LB. Differential Loss of Spinal Interneurons in a Mouse Model of ALS. Neuroscience. 2020;450:81–95. Epub 2020/08/29. doi: 10.1016/j.neuroscience.2020.08.011. PubMed PMID: 32858144.

