## Supplemental Figure 1 for "Hindlimb Muscle Representations in Mouse Motor Cortex Defined by Viral Tracing"

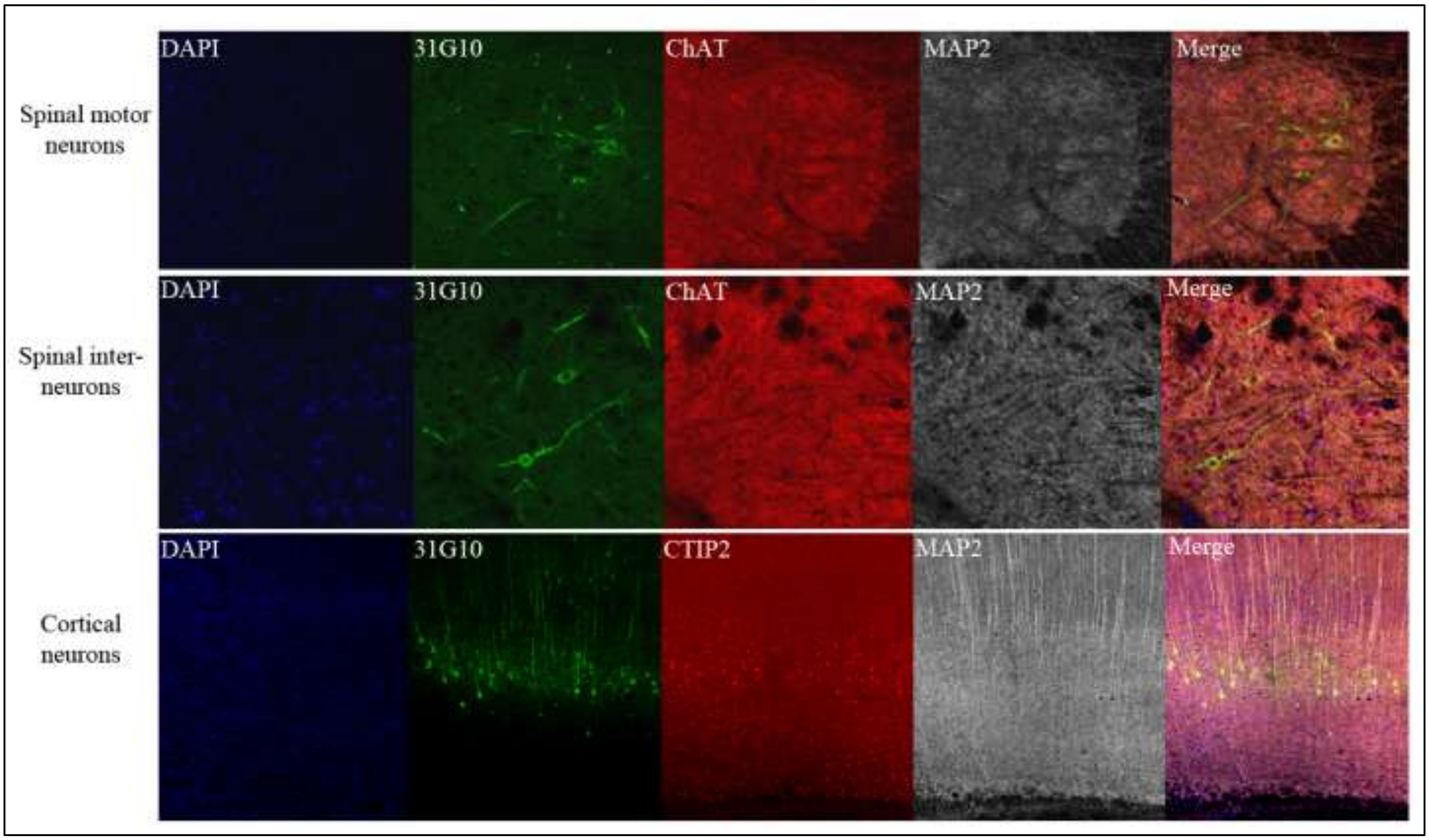


**Supplemental Figure 1**. Neurons located at specific anatomical nodes are infected in a time-dependent manner following rabies virus injection. Representative immunofluorescence images following viral injection into TA depict labeling in spinal motor neurons only ($31G10^{+}/ChAT^{+}/MAP2^{+}$; top row of panels), spinal interneurons ($31G10^{+}/ChAT^{-}/MAP2^{+}$; second row of panels), and cortical neurons ($31G10^{+}/CTIP2^{+}/MAP2^{+}$; third row of panels).
