## Supplemental Figure 3 for "Hindlimb Muscle Representations in Mouse Motor Cortex Defined by Viral Tracing"

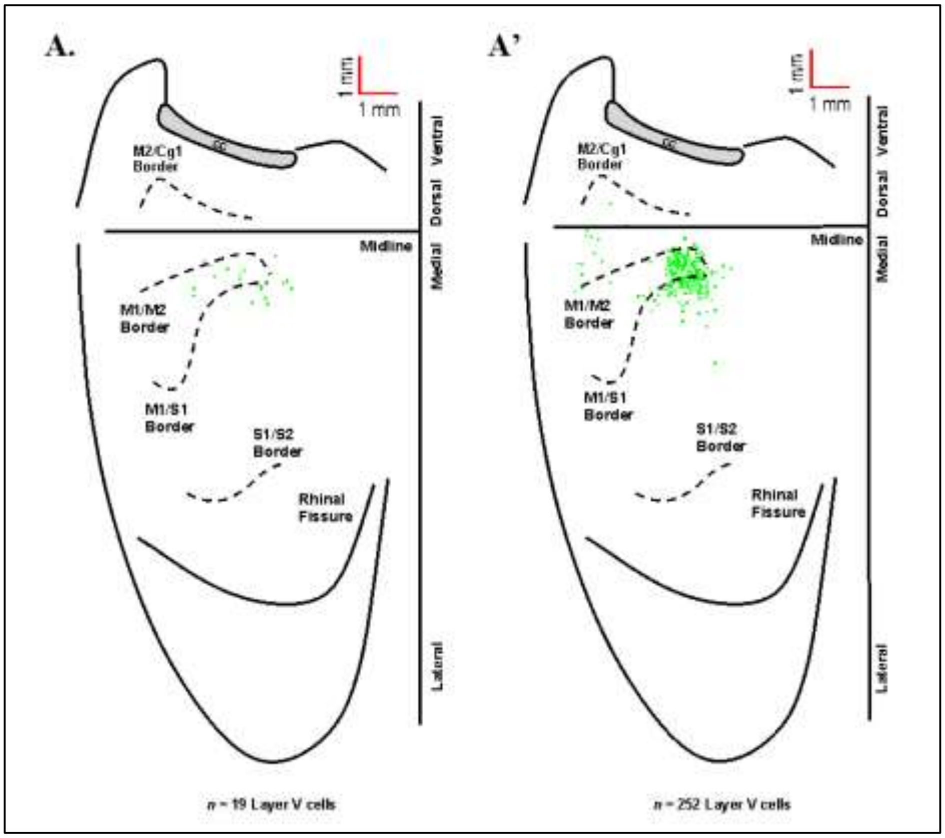


**Supplemental Figure 3**. Origin of cortical projections to soleus motor neurons. Maps of third-order neurons (square = one cell) in layer V that were labeled in the contralateral hemisphere after retrograde transneuronal transport of RV from the right soleus muscle [**A**: 60 hours (*n* = 4); **A'**: 67 hours (*n* = 4)]. Each map is a composite of multiple experiments that were overlapped on the atlas template. The medial wall of the contralateral hemisphere has been reflected upward and joined to the lateral surface at the midline. Cytoarchitectonic borders are delineated including the border between granular (S1) and agranular (M1) cortex in the region of the forelimb and hindlimb representations. Midline, midline of the hemisphere; M1, primary motor cortex; M2, secondary motor cortex (rostromedial motor field); S1, primary somatosensory cortex; S2, secondary somatosensory cortex; Cg1, cingulate cortex; CC, corpus callosum.
